# The activation of a *Suv39h1*-repressive antisense lncRNA by OCT4 couples the control of H3K9 methylation to pluripotency

**DOI:** 10.1101/2021.09.07.459323

**Authors:** Laure D. Bernard, Agnès Dubois, Victor Heurtier, Almira Chervova, Alexandra Tachtsidi, Noa Gil, Nick Owens, Sandrine Vandormael-Pournin, Igor Ulitsky, Michel Cohen-Tannoudji, Pablo Navarro

## Abstract

Histone H3 Lysine 9 (H3K9) methylation, a characteristic mark of heterochromatin, is progressively implemented during development to contribute to cell fate restriction as differentiation proceeds. For instance, in pluripotent mouse Embryonic Stem (ES) cells the global levels of H3K9 methylation are rather low and increase only upon differentiation. Conversely, H3K9 methylation represents an epigenetic barrier for reprogramming somatic cells back to pluripotency. How global H3K9 methylation levels are coupled with the acquisition and loss of pluripotency remains largely unknown. Here, we identify SUV39H1, a major H3K9 di- and tri-methylase, as an indirect target of the pluripotency network of Transcription Factors (TFs). We find that pluripotency TFs, principally OCT4, activate the expression of an uncharacterized antisense long non-coding RNA to *Suv39h1*, which we name *Suv39h1as*. In turn, *Suv39h1as* downregulates *Suv39h1* transcription in cis via a mechanism involving the modulation of the chromatin status of the locus. The targeted deletion of the *Suv39h1as* promoter region triggers increased SUV39H1 expression and H3K9me2 and H3K9me3 levels, leading to accelerated and more efficient commitment into differentiation. We report, therefore, a simple genetic circuitry coupling the global levels of H3K9 methylation to pluripotency in mouse ES cells.

## Introduction

During development, the establishment and maintenance of distinct gene expression patterns supporting the identity of each cell type are closely linked to the regulation of chromatin states^**1**^. Two broad states have been clearly and unambiguously identified: euchromatin, associated with transcriptionally active regions, and heterochromatin, associated with gene repression^**2-5**^. Two major states of heterochromatin have been traditionally considered. Facultative heterochromatin refers to a repressive chromatin environment displaying high variability across developmental stages, cell types and cell states. Indeed, silent developmental genes are usually embedded in facultative heterochromatin^**3,5**^. In contrast, ubiquitously silent elements such as retrotransposons and pericentromeric regions are locked by constitutive heterochromatin^**4,5**^. These two types of heterochromatin have been thought to be distinguishable by distinct molecular signatures, with facultative heterochromatin being characterized by trimethylation of histone H3 lysine 27 (H3K27me3) and constitutive heterochromatin by H3K9me3, among other chromatin features^**2-5**^. Nevertheless, recent data has challenged these strict definitions^**3**^. On the one hand, constitutive heterochromatin can under some circumstances be transcribed or decorated by marks previously associated with facultative heterochromatin^**6-8**^. On the other, while H3K27me3 and H3K9me2 were considered as major repressive mark for developmental genes, an increasing body of evidence points to H3K9me3 as an additional mean to silence developmental regulators as their expression is definitely shut down in particular lineages^**9**^. Hence, even though the role of H3K9 methylation in genome stability is unquestionable^**10**^, its importance in gene regulatory mechanisms during development appears to be equally important. Indeed, mouse knock-out (KO) models of H3K9 histone methyltransferases (HMTs) display penetrant phenotypes, particularly during gastrulation when pluripotency is lost and major differentiation events take place^**11,12**^. Conversely, before reaching pluripotency during early mouse embryogenesis the levels of H3K9 methylation are strictly controlled; promoting their increase, for instance by overexpressing the HMT SUV39H1, leads to developmental defects at the compaction stage^**13,14**^.

While extensive research has contributed to our understanding of how the establishment and maintenance of H3K27me3 regulates developmental transitions, how the levels of H3K9 methylation are developmentally regulated is less clear. Yet, a major distinction has been identified, particularly using pluripotent cells such as mouse Embryonic Stem (ES) cells. Indeed, H3K27me3 characterizes developmental genes even before differentiation, when they are embedded in the so-called bivalent chromatin, which is simultaneously enriched for H3K27me3 and for marks of activity^**15**^. Upon differentiation, H3K27me3 is either consolidated or erased in a cell-type-dependent manner^**16**^. On the contrary, H3K9 methylation is more largely controlled at the level of its abundance: during differentiation the global levels of H3K9me2 and H3K9me3 increase drastically^**17,18**^. Conversely, during the induction of pluripotency *in vitro* through reprogramming processes, H3K9 methylation has been shown to act as a major epigenetic barrier that is in part overcome by globally reducing its levels^**18,19**^. Therefore, while H3K27 methylation is mainly controlled by altering its genomic distribution, the global levels of H3K9 methylation display correlated changes to the differentiation status. Beyond the role of H3K9 methylation to stabilise somatic cell identities^**20**^, how its global levels are seemingly coupled to the acquisition and loss of pluripotency, and what consequences this coupling has, remain open questions.

In this study, we aimed at understanding the molecular basis of the link between H3K9 methylation and pluripotency. We find *Suv39h1* to be the only HMT tightly connected to the network of transcription factors (TFs) supporting pluripotency, particularly to its main player *Oct4*. The analysis of the mechanisms of *Suv39h1* repression by OCT4 led us to the identification of a repressive antisense long non-coding RNA (lncRNA^**20**^) to the *Suv39h1* gene. Our work identifies a simple genetic network based on the activation of the *Suv39h1* antisense by OCT4 which, in turn, represses *Suv39h1* transcription thereby coupling H3K9 methylation to pluripotency. Using CRISPR-Cas9 mediated deletion of the antisense promoters, we show that its activity controls the timing and efficiency of ES cell commitment into differentiation. Our results therefore provide a mechanistic perspective into how the global levels of H3K9 methylation are regulated at the onset of differentiation to irreversibly exit pluripotency.

## Results

### *Suv39h1* expression is under the control of OCT4 in ES cells

Using immunofluorescence, we first confirmed that differentiation of ES cells by LIF withdrawal leads to an increase of both H3K9me2 and H3K9me3 (**Fig.1A**; p<10^−15^), as expected. We hypothesized that one or several histone methyl-transferases or lysine demethylases (HMTs and KDMs, respectively)^**2,4**^ could be differentially expressed upon differentiation, linking the loss of pluripotency to increased H3K9 methylation. To assess this, we monitored mRNA levels of HMTs and KDMs using published RNA-seq datasets^**22**^ **(Fig.1B; Table S1)**. We found three HMTs to be upregulated upon differentiation: *Suv39h1* (p=0.004), *Suv39h2* (p=0.044) and *Glp* (p=0.004). In contrast, all tested KDM displayed minor changes below 2-fold. We reasoned that the increase of *Suv39h1, Suv39h2* and *Glp* expression could either be due to a direct control of their transcription by pluripotency TFs or to alternative mechanisms. To address this, we assessed the impact of the loss of individual pluripotency TFs (*Oct4, Nanog* and *Esrrb*) using available dox-inducible knock-outs^**23-25**^ **(Fig.S1A)**. Only one, *Suv39h1*, was found upregulated 24h after inducing the loss of pluripotency TFs, particularly of OCT4, which depletion leads to a 2-fold increase in *Suv39h1* mRNA levels **(Fig.1B; Table S1)**. Hence, after confirming *Suv39h1* expression changes by RT-qPCR **(Fig.S1B)**, we hypothesized that OCT4 may act as a repressor of *Suv39h1* expression to maintain low levels of H3K9 methylation until the onset of differentiation. Exploration of available ChIP-seq datasets^**26**^ **(Fig.1C)** and direct validation by ChIP-qPCR **(Fig.S1C)** identified a hotspot of pluripotency TFs, including OCT4, in the vicinity of *Suv39h1*. However, this TF binding hotspot was found located 3’ to *Suv39h1*, at around 27 kb of its promoter region. Notably, we noticed that this region coincides with the promoter region of an un-characterized gene, *Gm14820* (*AK010638*), antisense to and largely overlapping *Suv39h1* (top of **Fig.1C**).

**Fig. 1.**
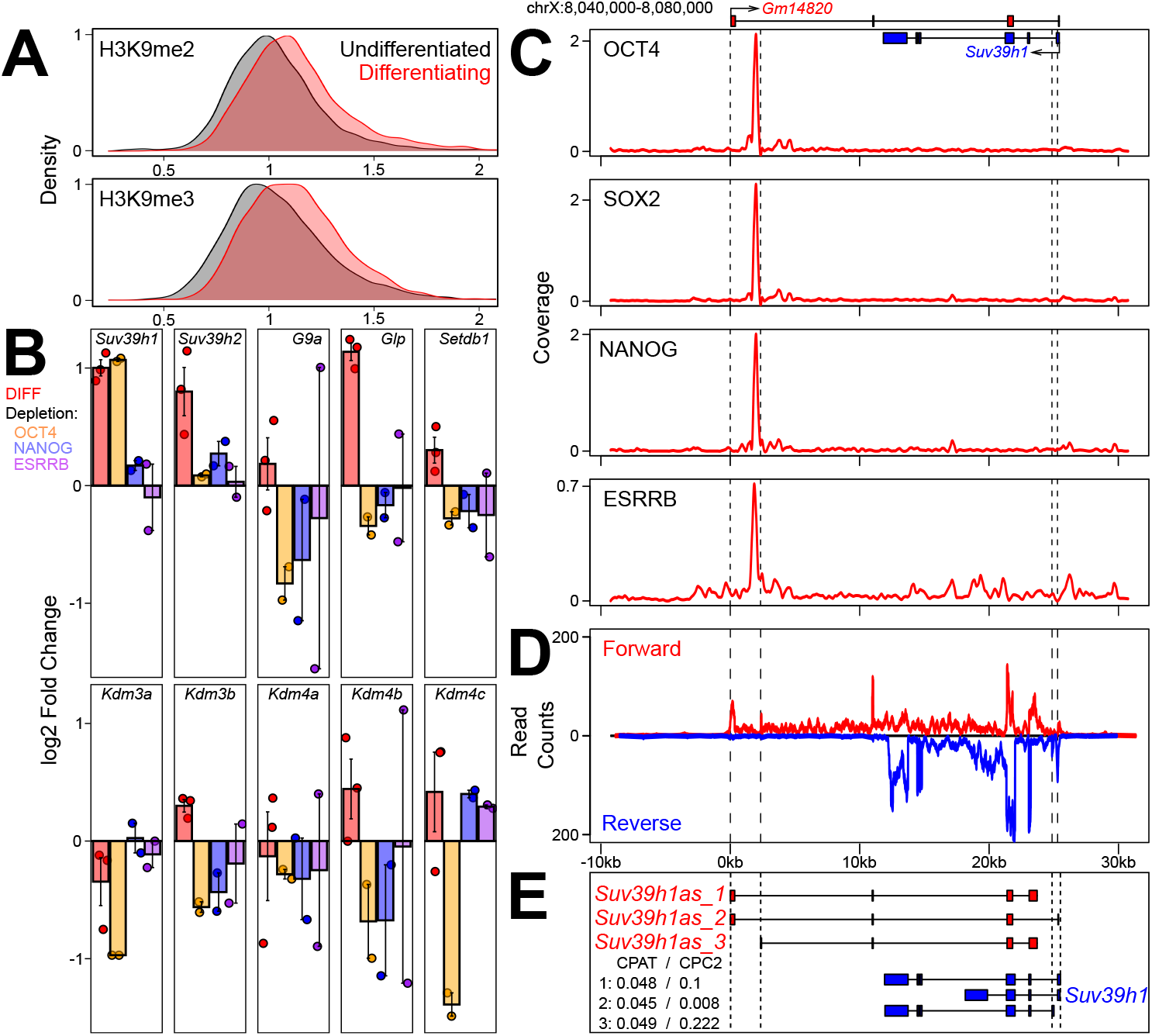
*Suv39h1* is downregulated by OCT4, which binds to the promoter of a *Suv39h1* antisense lncRNA. **(A)** Distribution of H3K9me2 (top) and H3K9me3 (bottom) in undifferentiated and differentiating ES cell populations assessed by immunofluorescence (Y-axis, density; X-axis, relative mean intensity per cell). Undifferentiated cells (black) are E14Tg2a cells cultured in FCS/LIF (black; n= 4503 cells for H3K9me2 and 3150 for H3K9me3); differentiating cells were obtained by LIF withdrawal (red; n= 6231 cells for H3K9me2 and 3755 cells for H3K9me3). **(B)** Log2 fold change of the indicated gene after differentiating ES cells as in (A) or 24h after inducing the depletion of individual TFs (OCT4, NANOG or ESRRB, as indicated) using Dox-inducible knock-out cells. Each dot represents an independent replicate and the bar and error bars the corresponding means and standard errors. **(C)** Average binding profile of OCT4, SOX2, NANOG and ESRRB (reads per million) across the *Suv39h1*/*Gm14820* locus (mm10, chrX:8,040,000-8,080,000 – 40 kb). *Suv39h1* and *Gm14820* are schematically represented on top. **(D)** RNA-seq profile across the *Suv39h1*/*Gm14820* locus, with forward and reverse fragment counts expressed with positive and negative values. **(E)** Schematic representation of *Gm14820*/*Suv39h1as* (red) and *Suv39h1* (blue) isoforms. The coding probabilities calculated with CPAT and CPC2 algorithms are shown for the three isoforms of *Suv39h1as*. The vertical dashed lines in (C), (D) and (E) mark the position of *Suv39h1as* or *Suv39h1* promoters.

### Identification of a *Suv39h1* antisense lncRNA

Stranded RNA-seq confirmed *Gm14820* to be expressed in ES cells, at levels comparable to *Suv39h1* **(Fig.1D)**. Using *de novo* transcript assembly with our RNA-seq datasets, together with direct cDNA cloning, sequencing and RT-qPCR, we identified several isoforms expressed in ES cells **(Fig.1E)**. All isoforms initiate from two distinct promoters, located in proximity to the region bound by pluripotency TFs, exhibit overlapping exons with *Suv39h1* and terminate within *Suv39h1* or in the vicinity of its 5’ end. Notably, *Gm14820* is annotated as a lncRNA^**21**^. Accordingly, using two different algorithms (CPAT^**27**^ and CPC2^**28**^), the nearly absent coding potential of all *Gm14820* isoforms was confirmed **(Fig.1E)**. Thereafter, we refer to *Gm14820* as *Suv39h1as*. To further characterize *Suv39h1as*, we assessed the stability of its RNA products and found the half-life of its spliced and unspliced forms to be around 12h and 1h30, respectively **(Fig.2A)**. However, *Suv39h1as* splicing is relatively inefficient compared to *Suv39h1* or another protein coding gene, *Nanog* **(Fig.2B)**, as is generally the case for lncRNAs^**21**^. Moreover, *Suv39h1as* was efficiently captured in poly-A selected RNA-seq, suggesting it is normally poly-adenylated **(Table S1)**. Next, we aimed at visualizing *Suv39h1as* RNA molecules in single cells. For this, we designed oligonucleotides targeting *Suv39h1as* exons and performed strand-specific single molecule RNA-FISH (smFISH) coupled to DNA-FISH to identify the *Suv39h1as*/*Suv39h1* locus, using a fosmid covering the whole region **(Fig.2C)**. We observed that *Suv39h1as* is mainly detected as a bright point in the nucleus, likely representing actively transcribed loci as it coincides with the DNA-FISH signal. A small number of single *Suv39h1as* RNA molecules could also be detected diffusing in the nucleus and, more rarely, in the cytoplasm. Quantification of the smFISH/DNA-FISH suggested a transcriptional frequency of around 50% in the population, with a median of 6 freely diffusing RNAs in cells presenting a transcriptionally active locus **(Fig.2D)**. Hence, we conclude that the pluripotency TFs bind close to the two promoters of a *Suv39h1* antisense lncRNA, which is mostly localised at its site of transcription, poly-adenylated and poorly spliced even though the spliced isoforms are relatively stable.

**Fig. 2.**
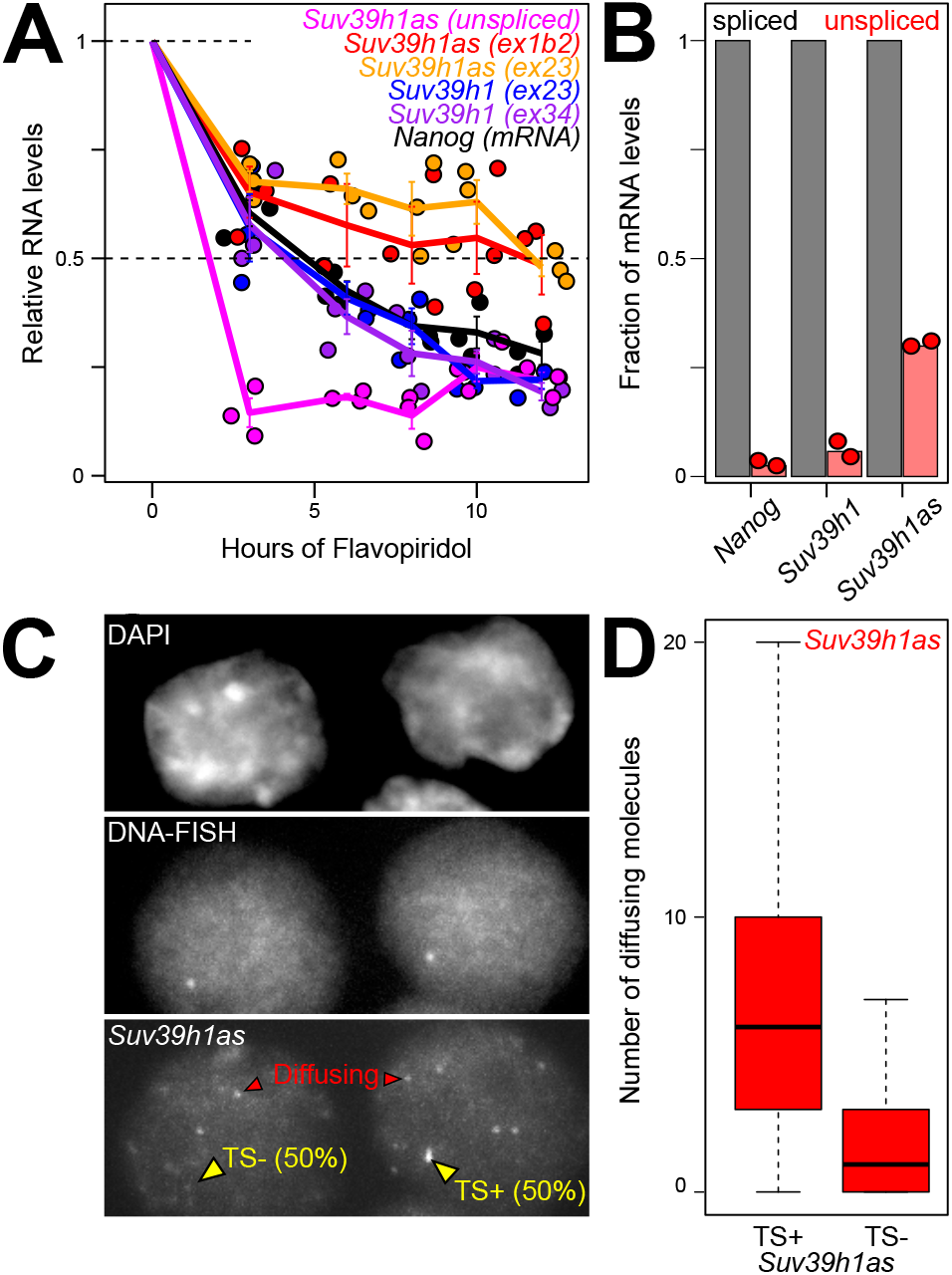
*Suv39h1as* is a nuclear, stable and lowly expressed lncRNA. **(A)** RT-qPCR analysis of the half-life of several RNA species during a transcription inhibition assay with Flavopiridol (X-axis, hours of treatment): *Suv39h1* mRNA, using two trans-exonic primer pairs between exons 2 and 3 (blue – ex23) or exons 3 and 4 (purple – ex34); *Suv39h1as*, using two trans-exonic primer pairs between exons 1b and 2 (red – ex1b2) or exons 2 and 3 (orange – ex23) or primer pairs amplifying the unspliced RNA (magenta – pre-mRNA); *Nanog* mRNA (black). Each dot represents an independent replicate and the line the corresponding mean and standard error. Ribosomal RNA (*28S*) was used for normalization. **(B)** Histogram representing unspliced RNA levels (red) relative to corresponding spliced RNAs (black) for *Nanog, Suv39h1* and *Suv39h1as*, as measured by RNA-seq. Each dot represents an independent replicate and the bar the corresponding mean. **(C)** Representative sm-FISH followed by DNA-FISH visualizing *Suv39h1as* RNA molecules and the *Suv39h1*/*Suv39h1as* locus, respectively, in undifferentiated E14Tg2a cells. Red arrowheads indicate RNAs diffusing away from the locus, which is indicated by a yellow arrow. The proportion of actively transcribing cells is indicated (n=358 cells). **(D)** Boxplots (median; 25-75% percentiles; error bars) showing the number of *Suv39h1as* diffusible molecules counted in cells presenting an active (TS+) or inactive (TS-) *Suv39h1as* gene (n=358).

### Anticorrelated expression patterns of *Suv39h1* and *Suv39h1as*

To investigate *Suv39h1as* and *Suv39h1* expression patterns we differentiated ES cells using three independent protocols based on LIF withdrawal, N2B27 or EpiLC-directed differentiation. These three assays showed a strong reduction of *Suv39h1as* expression after 3 days of differentiation, when *Suv39h1* expression increases (**Fig.3A**; p<0.05 (*Suv39h1*) and p<0.01 (*Suv39h1as*) for all differentiation assays). Moreover, the depletion of OCT4 showed a rapid downregulation of *Suv39h1as* (p<10^−3^ at all time-points), reaching minimal levels of expression within 12h and accompanied by a marked increase of *Suv39h1* expression (p<0.05 at all time-points) that reached maximal levels after 24h **(Fig.3B)**. Thus, *Suv39h1* and *Suv39h1as* display anticorrelated expression levels. To explore this observation at the single cell level, we designed oligonucleotides across *Suv39h1* exons and introns to monitor *Suv39h1*/*Suv39h1as* expression by smFISH in parallel to DNA-FISH **(Fig.3C,D)**. We found around 20% of cells actively transcribing both sense/antisense genes and around 30% transcribing either one or the other. Moreover, cells actively transcribing *Suv39h1as* displayed significantly (p<10^−11^) fewer *Suv39h1* mRNA molecules **(Fig.3D)**. In summary, both smFISH and RT-qPCR experiments showed that *Suv39h1* and *Suv39h1as* display anticorrelated expression patterns. This anticorrelation stems from single cell dynamics where the transcription of the antisense leads to a slight reduction of the transcriptional frequency of *Suv39h1* and, more significantly, to reduced expression of diffusing mRNAs. These observations indicate that *Suv39h1as* acts as a repressor of *Suv39h1*. Since *Suv39h1as* is downregulated upon differentiation and upon the loss of OCT4, our data suggests that pluripotent TFs activate *Suv39h1as* transcription which, in turn, downregulates *Suv39h1* expression.

**Fig. 3.**
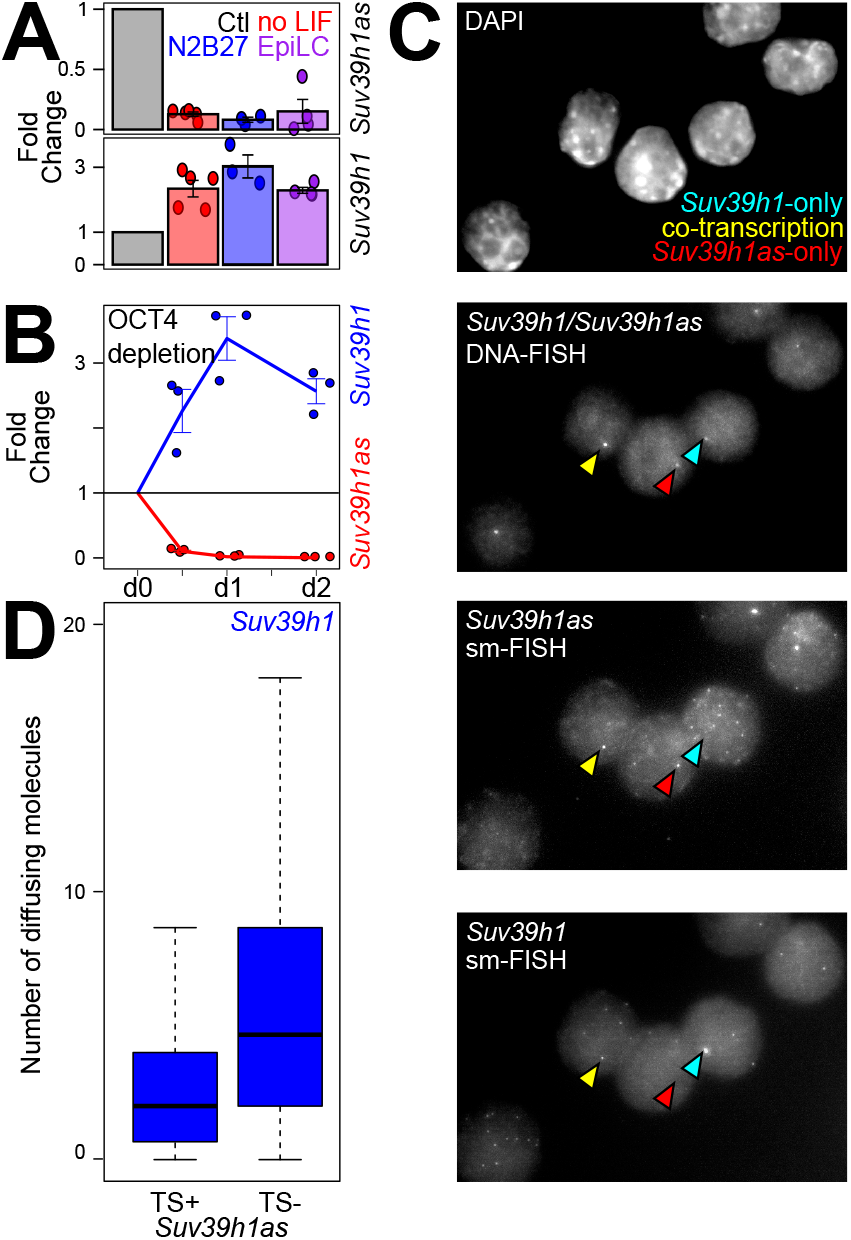
Anticorrelated expression of *Suv39h1* and *Suv39h1as*. **(A)** Fold change expression of *Suv39h1as* (top) or *Suv39h1* (bottom) measured by RT-qPCR in differentiating E14Tg2a cells versus undifferentiated controls (ctl; gray). Differentiation was triggered for three days with three independent protocols: LIF withdrawal from FCS/LIF cultures (no LIF; red), 2i and LIF withdrawal from 2i+LIF cultures (N2B27; blue) or EpiLC differentiation from 2i+LIF cultures (EpiLC; purple). Values were normalized to *Tbp* and fold changes calculated to their respective control culture. Each dot represents an independent replicate and the bar and error bars the corresponding means and standard errors. **(B)** RT-qPCR analysis of *Suv39h1as* (red) and *Suv39h1* (blue) expression upon OCT4 depletion for the indicated time (X-axis). Values were normalized to *Tbp*. Each dot represents an independent replicate and the line the corresponding mean with standard errors. **(C)** Representative sm-FISH of *Suv39h1as* and *Suv39h1* RNA molecules, followed by DNA-FISH visualising the *Suv39h1*/*Suv39h1as* locus in E14Tg2a cells (n= 358). Selected loci transcribing either *Suv39h1, Suv39h1as* or both genes are indicated with arrow heads: blue, *Suv39h1*-only (30%); red, *Suv39h1as*-only (30%); yellow for cells transcribing both (20%). **(D)** Boxplots (median; 25-75% percentiles; error bars) showing the number of *Suv39h1* diffusible molecules counted in cells presenting an active (TS+) or inactive (TS-) *Suv39h1as* gene, counted in 358 E14Tg2a cells.

### A simple OCT4-*Suv39h1as*-*Suv39h1* circuitry couples H3K9 methylation to pluripotency

To functionally establish the relationships between OCT4, *Suv39h1as, Suv39h1* and H3K9 methylation, we designed two gRNAs to delete 5,5kb encompassing the two promoters of *Suv39h1as*. Two independent KO clones, A8 and D8, were generated. PCR genotyping confirmed the deletion and analysis of their karyotypes did not reveal any overt defect. Moreover, RT-qPCR showed a complete loss of *Suv39h1as* expression **(Fig.4A)**. We then addressed the impact of *Suv39h1as* depletion on *Suv39h1* expression, both before and during differentiation. In the two mutant clones we observed an increase of *Suv39h1* expression in undifferentiated cells (p=0.011 and 0.0016 for A8 and D8, respectively), reaching the levels observed upon differentiation in WT cells **(Fig.4A)**. In differentiating cells, when *Suv39h1as* is naturally silenced, the deletion had no impact **(Fig.4A)**, as expected, regardless of the differentiation protocol **(Fig.S1D)**. Therefore, these results indicate that *Suv39h1as* acts as a pluripotency-associated repressor of *Suv39h1* expression. However, this does not imply that OCT4 exclusively represses *Suv39h1* expression via *Suv39h1as*. To address this, we used siRNAs targeting *Oct4* to test whether in the absence of *Suv39h1as*, the loss of OCT4 would lead to any modification of *Suv39h1* expression. Whereas in wild-type cells the knock-down of *Oct4* (above 80% efficiency, **Fig.S1E**), led to higher (p=0.027) *Suv39h1* expression **(Fig.4B)**, as expected, in mutant cells it was fully inconsequent **(Fig.4B)**. This indicates that the OCT4-dependent repression of *Suv39h1* is exclusively mediated by *Suv39h1as*. Next, we performed sm-FISH to study *Suv39h1* upregulation with single cell resolution **(Fig.4C)**. We observed a marked increase in the transcriptional frequency of *Suv39h1*, rising from 47.6% in WT to 76.3% and 74.1% in A8 and D8, respectively (p<10^−10^ for both clones). Moreover, the number of *Suv39h1* mRNAs per cell also increased substantially (p<10^−15^), with virtually no cell displaying an absence of *Suv39h1* mRNAs **(Fig.4D)**. This increase in *Suv39h1* expression was accompanied by higher levels of SUV39H1 protein expression and accumulation at heterochromatic regions such as chromocenters, as established by immuno-staining and western-blot **(Fig.4E,F; Fig.S1F)**. Consequently, in both mutant clones we observed higher levels of H3K9me2 and H3K9me3 (p<10^−8^ for both marks and cell clones), establishing a direct link between *Suv39h1as* and the global levels of H3K9 methylation in ES cells **(Fig.4G)**.

**Fig. 4.**
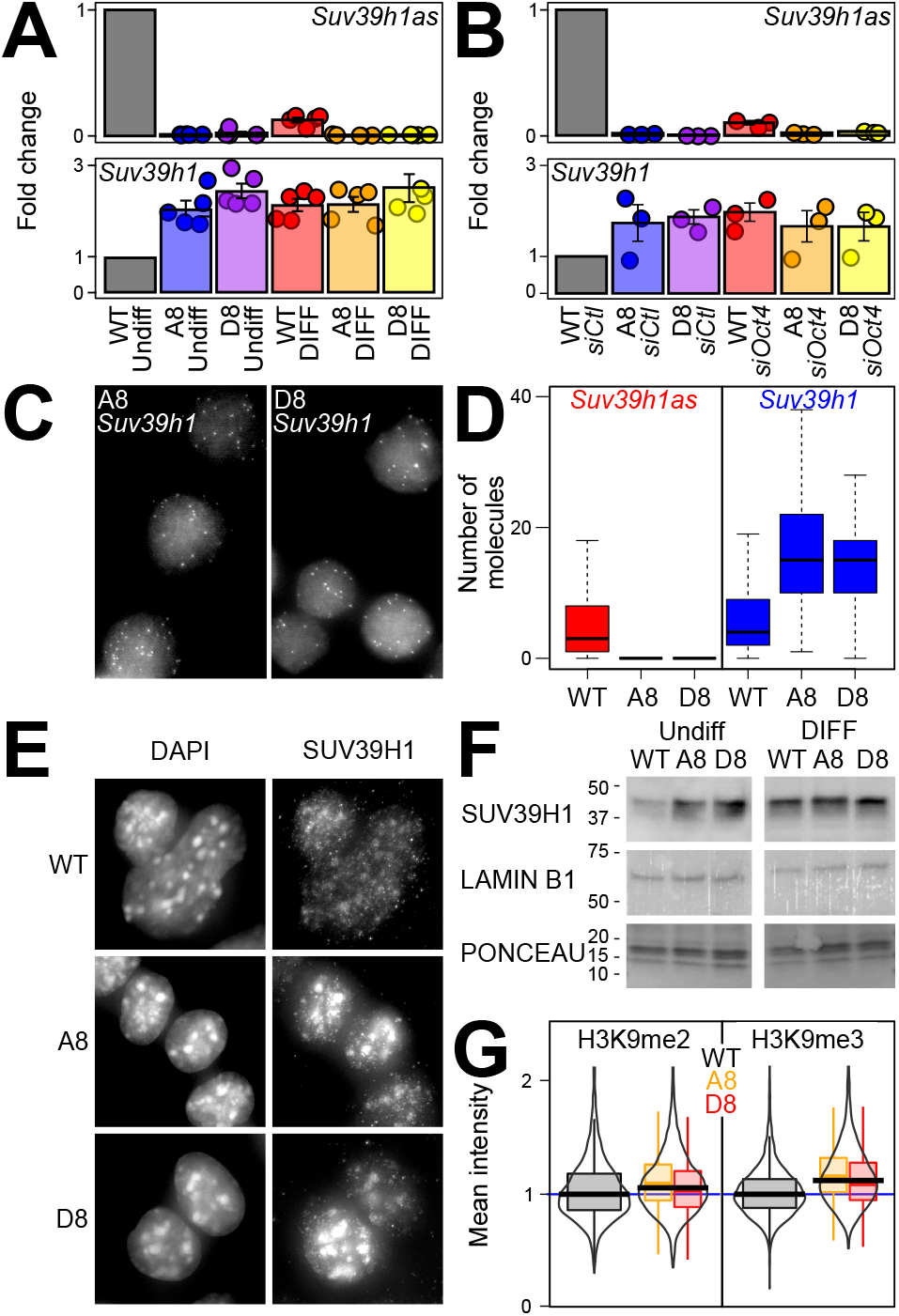
OCT4 represses *Suv39h1* via *Suv39h1as* to reduce H3K9 methylation in ES cells. **(A)** Expression fold change of *Suv39h1as* (top) and *Suv39h1* (bottom) in WT (E14Tg2a) and *Suv39h1as*-mutant cells (A8 and D8) cultured in un-differentiated (FCS/LIF) or differentiating conditions (3 days without LIF). Values were normalized to *Tbp*. Each dot represents an independent replicate and the bar and error bars the corresponding means and standard errors. **(B)** Expression fold change of *Suv39h1as* (top) and *Suv39h1* (bottom) in wild-type (WT – E14Tg2a) and *Suv39h1as*-mutant cells (A8 and D8) knocked-down with either control or *Oct4*-targetted siRNAs. Values were normalized to *Tbp*. Each dot represents an independent replicate and the bar and error bars the corresponding means and standard errors. **(C)** Representative sm-FISH images of *Suv39h1* in A8 and D8 cells (wild-type cells presented in Fig.3C). **(D)** Boxplots (median; 25-75% percentiles; error bars) showing the number of *Suv39h1as* (red) or *Suv39h1* (blue) diffusible molecules counted in wild-type (WT; n= 358) or mutant cells (A8, n= 289; D8, n= 270). **(E)** Representative SUV39H1 immunofluorescence of wild-type (E14Tg2a) and *Suv39h1*-mutant ES cells (A8 and D8). **(F)** Representative Western-Blot of SUV39H1, LAMIN B1 and corresponding Ponceau for wild-type (E14Tg2) and mutant cells (A8 and D8) in undifferentiated (FCS/L) and differentiating (3 days without LIF) conditions. On the left of each image is indicated protein scale in kDa. **(G)** Violin and boxplots (median; 25-75% percentiles; error bars) presenting relative H3K9me2 (left) and H3K9me3 (right) mean intensity values of E14Tg2a (WT – black; n=12881 for H3K9me2 and n=12053 for H3K9me3) or mutant cells (A8, yellow, n=3553 cells for H3K9me2 and 2081 for H3K9me3; D8, red, n=5050 cells for H3K9me2 and 2641 for H3K9me3), assessed by immunofluorescence.

### *Suv39h1as* modifies the chromatin of the *Suv39h1as*/*Suv39h1* locus

We have observed that in mutant ES cells lacking *Suv39h1as* expression, the transcriptional frequency of *Suv39h1* increases from 50 to 75%. Moreover, the absence of *Suv39h1as* is not accompanied by increased stability of *Suv39h1* mRNAs **(Fig.S1G)**. Therefore, *Suv39h1as* is likely to act as a transcriptional repressor of *Suv39h1*. To explore this, and given that other antisense transcription units have been shown to modify the chromatin of their corresponding sense gene^**29-31**^, we used a ChIP approach to establish the histone modification profiles of the locus **(Fig.5)**. First, we monitored H3K4 methylation profiles. We found H3K4me1 and me2, which usually mark transcriptionally competent regions^**32**^, to globally decorate the locus with minimal focal accumulation at the promoters. Conversely, H3K4me3, a mark of activity^**32**^, was focally enriched at the *Suv39h1* promoter and displayed low levels over the antisense promoter. We then profiled the active histone acetylation marks, H3K9ac and H3K27ac. Similarly to H3K4me3, we found H3K9ac to preferentially mark the *Suv39h1* promoter. In contrast, both sense and antisense gene promoters where enriched for H3K27ac. In mutant cells we observed a global decrease of H3K4me1/me2 over the region transcribed by *Suv39h1as* (p<10^−3^), particularly before it reaches the *Suv39h1* gene body **(Fig.5)**, indicating its transcription promotes the establishment of these marks. The lack of H3K4me1/me2 reduction within the region transcribed by both genes suggests that the increased transcription of *Suv39h1* may have a compensatory role. Moreover, H3K4me2, H3K9ac and H3K27ac, all marks of gene activity, showed a slight but statistically significant increase (p<0.05) at *Suv39h1* promoter in the absence of *Suv39h1as* **(Fig.5)**. Altogether, this analysis suggests that the loss of *Suv39h1as* leads to increased *Suv39h1* transcription at least in part mediated by increased euchromatinisation of the Suvar39h1 promoter. These results suggest a simple mechanism where *Suv39h1as* modulates the local chromatin environment of *Suv39h1* to decrease the probability of its transcription, leading to a global lower level of *Suv39h1* mRNA and, subsequently, protein.

**Fig. 5.**
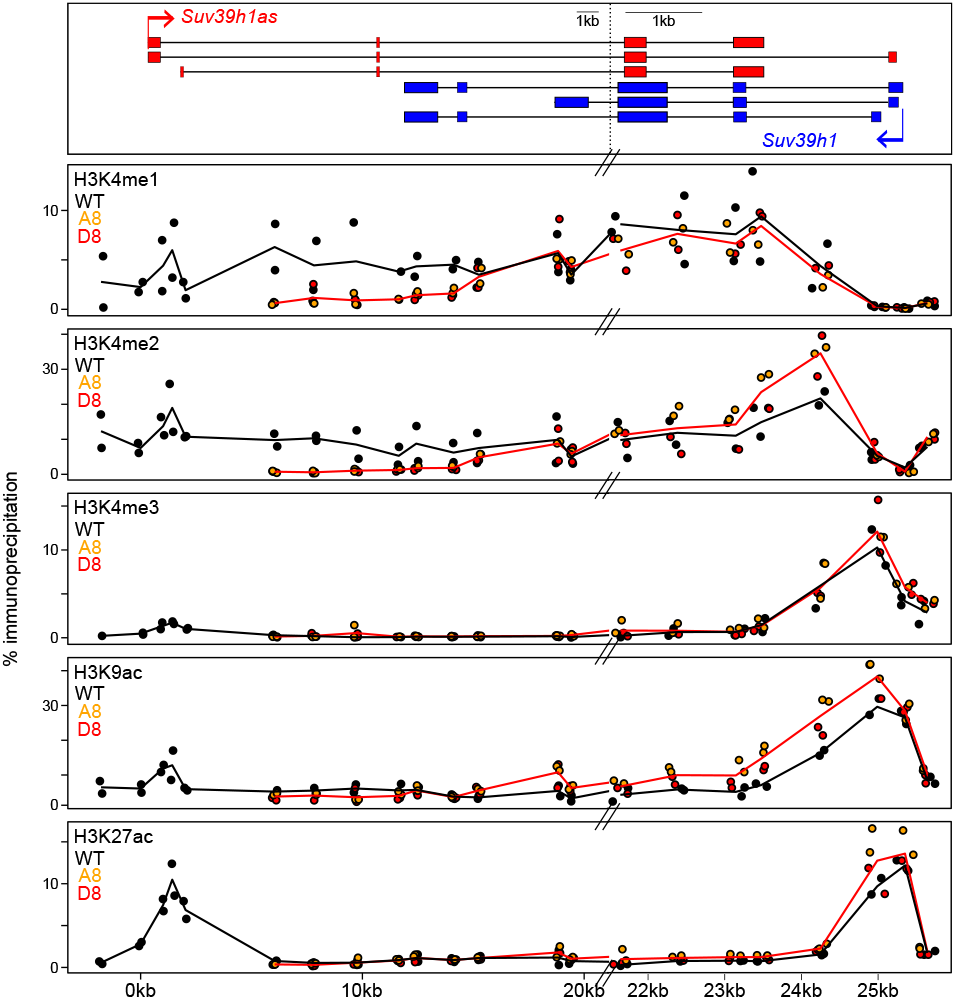
*Suv39h1as* triggers complex chromatin changes across the locus. Chromatin immunoprecipitation profile (Y-axis, percentage of immunoprecipitation) of H3K4me1, H3K4me2, H3K4me3, H3K9ac and H3K27ac, as indicated, across *Suv39h1*/*Suv39h1as* locus in wild-type (E14Tg2a, black) and *Suv39h1as*-mutant cells (A8, yellow dots; D8, red dots; the red line represents the average of all data points for mutant clones). The X-axis represents genomic distances in kb with respect to the *Suv39h1as* transcription start site, as schematized on top. Note a break on the scale of the genomic coordinates at around X=21kb.

### The lack of *Suv39h1as* leads to accelerated differentiation commitment

We finally wondered whether the increase of H3K9 methylation taking place in *Suv39h1as* mutant cells had any physiological impact. First, we used clonal assays to assess self-renewal and differentiation efficiency **(Fig.6A,B)**. Either in conditions of reinforced self-renewal (2i/LIF), in traditional serum-containing culture medium (FCS/LIF) or in the absence of LIF (FCS), the number of alkaline-phosphatase colonies, a marker of pluripotent cells, was identical between wild-type and mutant clones. Hence, the presence of increased H3K9 methylation is largely inconsequential for self-renewal and for the loss of pluripotency. In agreement, both wild-type and mutant cells proliferate and differentiate normally, as evaluated morphologically **(Fig.S1H)** and by marker expression **(Fig.S1I)**. However, during differentiation, the role of H3K9 methylation is to restrict cell fate and developmental competence^**18,33**^, more than to elicit differentiation. Therefore, we reasoned that the loss of *Suv39h1as* could modulate the timing of commitment into differentiation. To test this, we used an established assay^**34**^ whereby wild-type and mutant clones were differentiated in parallel and, every day, the cells were harvested and reseeded clonally in 2i/LIF: only those cells that were not yet committed into irreversible differentiation can self-renew and form undifferentiated colonies **(Fig.6C,D)**. As previously shown, we observed that commitment took place between days 2 and 3 in wild-type cells, with a reduction in clonogenicity of nearly 90% **(Fig.6D)**. In mutant cells, however, the reduction in the number of undifferentiated colonies was more marked from day 2.5 onwards (**Fig.6D**; p=0.0666, 0.0020 and 0.0041 for days 2.5, 3 and 4, respectively). Therefore, the premature establishment of higher levels of H3K9me2/me3 in ES cells facilitates the irreversible commitment into differentiation, in line with the role of these repressive marks in locking cell fate changes^**20**^.

**Fig. 6.**
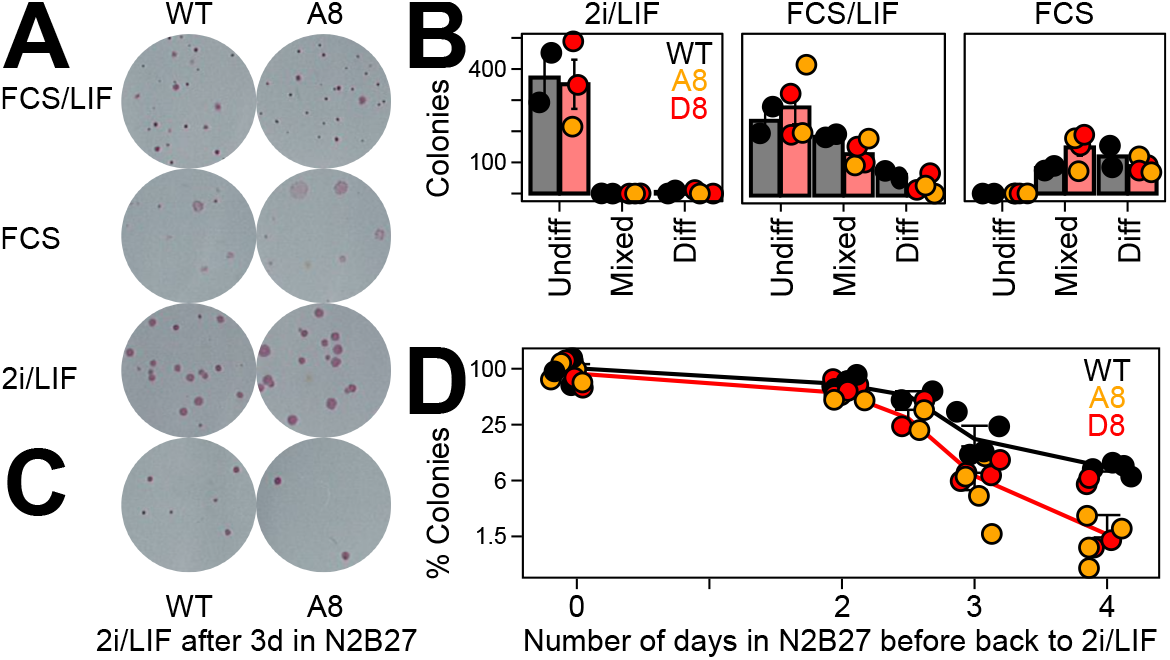
Accelerated commitment into differentiation in the absence of *Suv39h1as*. **(A)** Representative alkaline-phosphatase staining of ES cell colonies cultured as indicated. **(B)** Number of wild-type (E14Tg2a, black) or *Suv39h1as*-mutant (A8, orange points; D8, red points) colonies characterized as undifferentiated, mixed or differentiated after culturing them as indicated. Each dot represents an independent replicate and the bar and error bars the corresponding means (black for wild-type and red for the mean of all data points for mutant clones) and standard errors. **(C)** Alkaline-phosphatase staining of ES cell colonies cultured in 2i/LIF after 3 days in N2B27 for wild-type (E14Tg2a) and *Suv39h1as*-mutant cells (A8). **(D)** Percentage of alkaline-phosphatase positive colonies cultured in 2i/LIF after differentiating them for the indicated number of days (X-axis) in N2B27. D0, undifferentiated cells were set as 100%. Each dot represents an independent replicate (wild-type, black; *Suv39h1as*-mutant clones in orange, A8, and red, D8) and the line the corresponding mean and standard error (all mutant datapoints were averaged to obtain the red line).

## Discussion

In this study, we have identified a genetic network linking the control of the global levels of H3K9 methylation to pluripotency. The pluripotency network, mainly through OCT4, activates an antisense lncRNA to the *Suv39h1* gene (*Suv39h1as*), repressing *Suv39h1* expression. Consequently, the level of H3K9 methylation is reduced. Upon differentiation, the collapse of the pluripotency network leads to the silencing on *Suv39h1as*, enabling increased SUV39H1 expression and H3K9 methylation, which controls the timing and efficiency of the irreversible commitment into differentiation. This genetic setup may act as a time-delay generator, enabling ES cells to filter out transient fluctuations in the activity of the pluripotency network^**35**^: only a long decrease in activity of the pluripotency network may be sufficient to elicit the increase in SUV39H1 expression that will follow the extinction of *Suv39h1as*.

Antisense lncRNAs are frequent in mammals, with 29% of canonical protein coding genes displaying antisense transcription^**36**^. Given their antisense orientation and the resulting complementarity, antisense lncRNAs can theoretically regulate their cis-linked sense gene through a wide variety of mechanisms. By deleting the *Suv39h1as* promoter region, we found that *Suv39h1as* controls the frequency of *Suv39h1* transcription and has no impact on *Suv39h1* mRNA stability. However, the exact mechanisms by which *Suv39h1as* represses *Suv39h1* transcription remain open, even though we show it has complex influences on the chromatin. On the one hand, *Suv39h1as* triggers H3K4me1 and me2 throughout the locus; on the other, it slightly reduces H3K4me2, H3K4me3, H3K9ac and H3K27ac at the *Suv39h1* promoter. Notably, these effects are reminiscent of those associated with the regulation of *Xist* by its antisense *Tsix*^**29-31**^, possibly revealing a general property of antisense transcription. While it is possible that the slight increase of euchromatic marks at the *Suv39h1* promoter are sufficient to increase the frequency of *Suv39h1* transcription, other potential mechanisms cannot be excluded. For instance, the antisense/sense polymerases could enter into physical collision and transcriptional abortion^**37**^, modulating *Suv39h1* elongation. Moreover, *Suv39h1as* presents two exons nearly fully overlapping *Suv39h1* exons, a gene structure that is conserved at the human *Suv39h1as*/*Suv39h1* locus **(Fig.S1J)**. Therefore, while the transcriptional effects are clear, *Suv39h1as* may deploy additional post-transcriptional mechanisms to downregulate *Suv39h1* mRNA expression, such as masking splice sites as it has been reported for *ErbAa*^**38**^. Finally, other mechanisms fully independent of *Suv39h1as* transcription or of the produced lncRNA cannot be excluded, such as direct competition between the sense and antisense promoters^**39**^.

The deletion of *Suv39h1as* promoter and the ensuing increase in H3K9 methylation, appears to be largely inconsequent for ES cells: they self-renew and differentiate efficiently. This observation is in line with others, where histone modifiers have been either invalidated or ectopically expressed in ES cells with minor consequences for self-renewal^**40,41**^. However, despite the fact that *Suv39h1as* mutant cells self-renew and differentiate normally, we asked whether the timing of commitment into differentiation is altered. Our results showed a faster and more efficient commitment into differentiation, suggesting that the global levels of H3K9 methylation contribute to irreversibly lock the loss of pluripotency. This observation adds to the notion of H3K9 methylation acting as an epigenetic barrier providing robustness to cell fate changes^**33**^. Moreover, our results also underscore the dominance of pluripotency TFs over chromatin modifications^**41**^. We had already shown that NANOG, another key pluripotency TF, controls H3K27me3 levels, particularly during early differentiation^**22**^. Here, we complement this notion with OCT4 controlling H3K9me3 via the *Suv39h1as*/*Suv39h1* tandem. Together, these results place the control of global levels of heterochromatin marks under the activity of the pluripotency network, extending the concept of the genetic dominance of pluripotency. Whether our observations and conclusions can be extended to early mouse embryogenesis and to the acquisition and loss of pluripotency *in vivo* is now a question of primary importance. Notably, H3K9 methylation levels are exquisitely regulated during early embryogenesis^**42**^. It is noteworthy that SUV39H1 is absent in oocytes and its expression starts at the 2-4 cell transition stage^**43**^, when the reconfiguration of constitutive heterochromatin as chromocenters is initiated. Moreover, the overexpression of SUV39H1 during the early stages of embryogenesis leads to developmental defects. Indeed, using different strategies, Zhang et al.^**13**^ and Burton et al.^**14**^ over-expressed SUV39H1 in the zygote and observed an increase of H3K9me3 levels leading to early developmental arrest at the time of compaction. Therefore, an important hypothesis emerges from this work: *Suv39h1as* could be a key regulator of *Suv39h1* during early embryogenesis, holding its expression until the appropriate time to enable the timely establishing of the first heterochromatic structures in the embryo.

## Supplementary information

One supplementary figure accompanies this manuscript, it can be found at the end of this document. Two Supplementary Tables and Methods are available online.

## Acknowledgements

This study was conceived by P.N with inputs from L.B. and V.H.. Experiments were designed and executed by L.B., with help from A.D., V.H., A.T., S.V-P. and M.C-T. RNA-seq analyses and assembly were done by N.G., N.O., A.C. and I.U. The paper was written by L.B. and P.N. L.B. acknowledges the Ecole Normale Supérieure and Sorbonne Université for funding. P.N. and M.C-T. acknowledge the Labex Revive (Investissement d’Avenir; ANR-10-LABX-73), the Institut Pasteur and the CNRS for funding.

---

**Supplementary Information, Fig. S 1.**
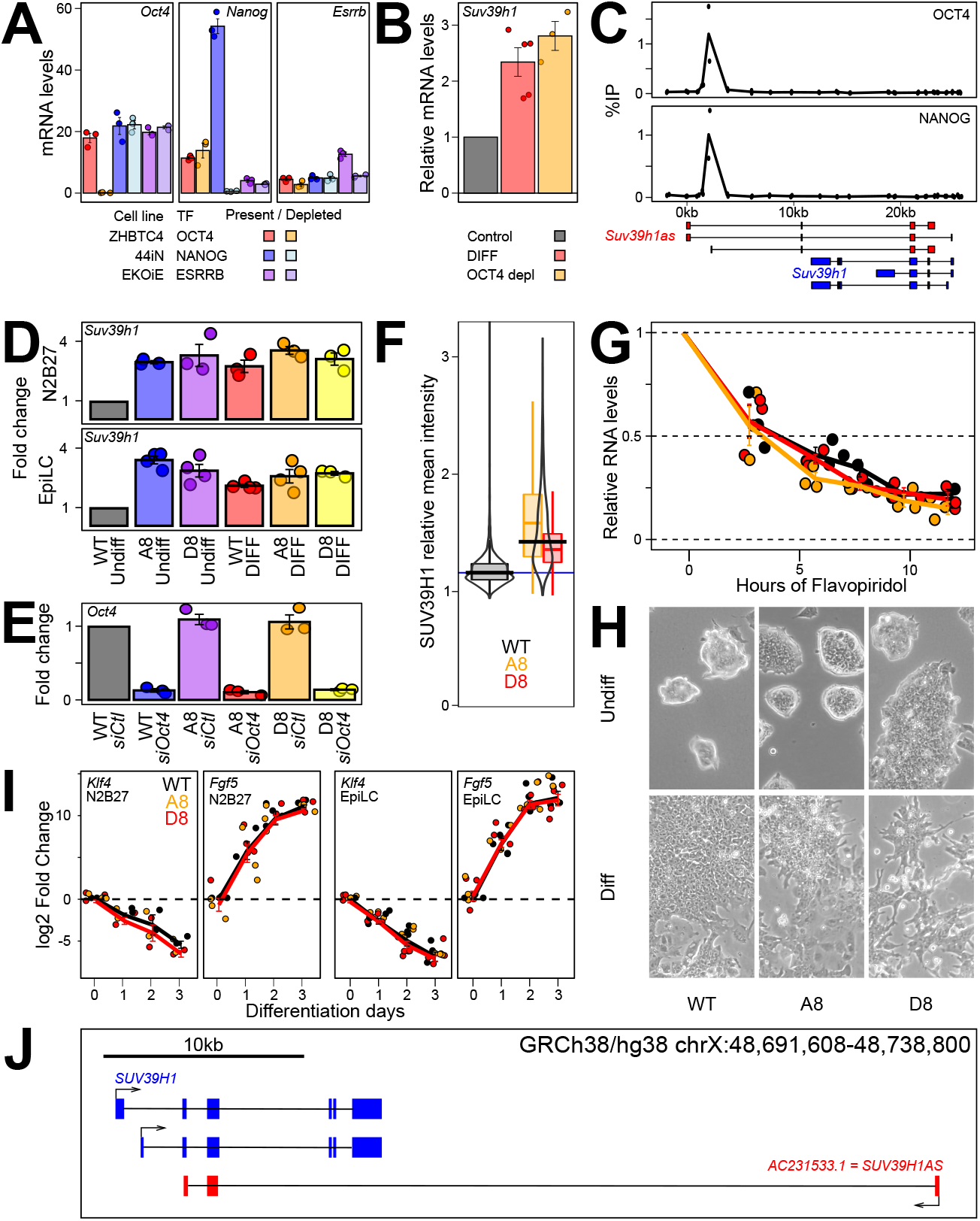
Additional information on *Suv39h1* regulation by OCT4 and *Suv39h1as*. **(A)** Expression of *Oct4, Nanog* or *Esrrb* upon inducing their depletion in specific dox-inducible knock-out lines. Note in EKOiE the remnant expression of *Esrrb* produces a truncated, non-functional protein^**23**^. Each dot represents an independent replicate and the bar the corresponding mean and standard error. **(B)** RT-qPCR validation of *Suv39h1* overexpression upon differentiation (DIFF, 3 days without LIF, red) or upon OCT4 depletion (24h, orange). Each dot represents an independent replicate and the histogram the corresponding mean and standard error. **(C)** ChIP-qPCR validation of OCT4 and NANOG binding at the promoter region of *Suv39h1as* in E14Tg2a (X-axis represents genomic distances in kb with respect to the *Suv39h1as* transcription start site). Each black dot represents the percentage of immunoprecipitation (%IP; Y-axis) of an independent replicate and the line the corresponding mean. A schematic representation of the locus with different isoforms is shown below. **(D)** RT-qPCR confirmation of lack of increased *Suv39h1* upregulation in differentiated *Suv39h1as*-mutant cells obtained by N2B27 or EpiLC assays as compared to wild-type cells. Each dot represents an independent replicate and the histogram the corresponding mean and standard error. **(E)** RT-qPCR confirmation of *Oct4* knock-down in wild-type and *Suv39h1as*-mutant cells. Each dot represents an independent replicate and the histogram the corresponding mean and standard error. **(F)** Violin and box-plots (median; 25-75% percentiles; error bars) showing immunofluorescence quantification of SUV39H1 mean intensity in wild-type (black; n= 4048 cells) and mutant cells (A8 – yellow; n= 4949 cells and D8 – red; n= 4448 cells). **(G)** Analysis of *Suv39h1* mRNA half-life in wild-type and *Suv39h1as*-mutant cells, performed and presented as in Fig. 2A. **(H)** Representative bright-field photomicrographs of wild-type (E14Tg2a) and mutant (A8 and D8) cells cultured in undifferentiating (FCS/LIF) and differentiating (3 days of LIF withdrawal) conditions. **(I)** RT-qPCR analysis of ES (*Klf4*) and differentiation (*Fgf5*) markers during differentiation in N2B27 or in EpiLC conditions, as indicated, in wild-type (E14Tg2A, black) and in *Suv39h1as*-mutant cells (A8, orange, and D8, red). Each dot represents an independent replicate and the lines corresponding means (Black for WT and red for both mutant) with error bars. **(J)** Schematic representation of the *SUV39H1/SUV39H1as* locus in the human genome (Gencode V36 assembly).

