## Supplementary material for "The activation of a *Suv39h1*-repressive antisense lncRNA by OCT4 couples the control of H3K9 methylation to pluripotency": Methods

**Supplementary Methods**

### OCT4 activates a *Suv39h1*-repressive antisense lncRNA to couple histone H3 Lysine 9 methylation to pluripotency

**Laure D. Bernard et al.**

*Cell lines and generation of A8 and D8 Suv39h1as mutant cells.*

Wild-type cells in this study are E14Tg2a ES cells, from which all mutant cells were derived. Dox-inducible knock-out cells have been described before (Esrrb: EKOiE[23], Oct4: Zhbtc4[24] and Nanog: 44iN[25]). To generate ES cells deleted for the Suv39h1as promoter, several gRNAs were designed using several online resources (sam.genome-engineering.org, crispr.mit.edu, chopchop.cbu.uib.no). Highly ranked gRNAs for off-targets as well as on-target predictions score were selected. The gRNAs were cloned into plasmids containing a U6-promoter driven expression cassette along with a puromycin selection cassette or the Cas9 enzyme linked to a mCherry reporter (addgene 51133 and 64324 respectively, see gRNA sequences on Table S2). E14Tg2a cells were lipofected (Lipofectamin 2000; Invitrogen, 11668-019) with 1 µg of the plasmid containing the right deletion site guide and the Cas9 cassette and 3 µg of the plasmid containing the left deletion site guide and a Puromycin cassette. Puromycin (1 µg/ml – Sigma, P9620-10ml) selection was performed for 3 days and mCherry fluorescence checked by microscopy. Resistant cells were plated at clonal density and colonies picked and expanded. Genomic DNA was isolated with NucleoSpin Tissue DNA extraction Kit (Macherey-Nagel, 740952.50), and screened by qPCR and PCR. PCR was performed with LongAmp Taq PCR kit (BioLabs, E5200S) following manufactory's instructions and sequenced. qPCR and PCR primers are available in Table S2. Two knock-out clones, A8 and D8, were selected for further analyses of *Suv39h1* expression. The selected clones were checked by PCR on genomic DNA with primers indicated on Table S2 and with LongAmp Taq PCR kit (BioLabs, E5200S) or Q5 High Fidelity (NEB, M0491S) following manufactory's instructions. The PCR product was validated by agarose gel migration and purified (NucleoSpin Gel and PCR Clean-Up; Macherey-Nagel, 740609.50). PCR products were purified and cloned (ThermoFisher, cat n°450245) as indicated by the manufacturer. Ten bacterial colonies were picked, expanded and checked by PCR and sequencing. The precisely expected deletion was observed for A8 (5.5kb); D8 showed a 5.7kb deletion exhibiting a shift of 700bp compared to A8 but encompassing both *Suv39h1as* promoters. Both clones were karyotyped, with A8 exhibiting a normal karyotype and D8 presenting 50% of cells with an extra chromosome.

*Regular cell culture.*

Cells were cultured at 37°C, 7% CO2 on 0.1% gelatin-coated plates (SIGMA, G1890-100G) in DMEM+GlutaMax-I (Gibco, 31966-021), 10% FCS (Sigma F7524), 100µM 2-mercaptoethanol (Gibco, 31350-010), 1X MEM non-essential amino acids (Gibco, 1140-035)], supplemented with 10 ng/ml recombinant Leukemia Inhibitory Factor (MILTENYI BIOTEC, 130-099-895). In 2i/LIF medium, cells were grown in N2B27 base [0.5X DMEM/F-12 (Life Technologies, 31331-093), 0.5X Neurobasal (Life Technologies, 21103- 049), 0.5X N2 (Life Technologies, 17502-048), 0.5X B27 (Life Technologies, 17504-044), Insulin 10 µg/ml (Sigma, I-1882), 2mM L-Glutamin (Life Technologies, 25030-024), 37.5 µg/ml BSA (Sigma, A3311-10G) and 0.1mM 2-mercaptoethanol (Life Technologies, 31350010)], supplemented with 10ng/ml recombinant LIF (MILTENY BIOTEC, 130-099-895), 1µM PD0325901 (Axon 1408) and 3µM CHIRON99021 (Axon 1386). Doxycycline (1000 ng/µL – D3072, Sigma), Puromycin (1 µg/ml – Sigma, P9620-10ml) or Flavopiridol (400 nM – Selleckchem, S2679), were extemporaneously added, as indicated. Cells were passaged every 2-3 days, when they reached 70-80% confluency.

*Differentiation of ES cells.*

Cells cultured in FCS/LIF were differentiated by seeding 300000 cells per well of gelatin-coated 6-wells plate and withdrawing LIF. N2B27 and EpiLC differentiation assays were performed with cells cultured in 2i/LIF for a minimum of 3 passages (9 days). For N2B27 differentiation, 50000 cells per well were seeded on wells of 6-wells plates coated overnight with poly-L-ornithine 0.01% (Sigma, Cat# P4957) at 37°C and 2h with 1X laminin (Sigma, Cat# L2020) and LIF, PD0325901 and CHIRON99021 were withdrawn. For EpiLC differentiation, 230000 cells were seeded per well of 6 wells plates coated with human plasma fibronectin (10 µg/well – F2006, Sigma) and cultured in N2B27 medium containing activin A (20 ng/ml – 338-AC-010, R&D Systems), rhFGF (12 ng/ml – 233-FB, R&Dsystems), and KSR (1% – 10-828-010, Gibco).

*Clonal and differentiation commitment assays.*

For comparing self-renewal and differentiation capacity of wild-type and *Suv39h1as* clones, 600 cells were plated in gelatin-coated wells of a six well plate. Cells were cultured for 7 days in the indicated media and stained for alkaline phosphatase activity (Sigma, cat. 86R-1KT), following the manufacturer's instructions. Colonies were scored as undifferentiated, mixed and differentiated using a stereo-microscope (NIKON-SMZ1500). For commitment assays, 600 cells obtained every day of differentiation in N2B27 were plated in poly-L-ornithine/laminin-coated wells of a 6 well plate, cultured for 7 days in 2i+LIF and stained for alkaline phosphatase activity.

*Assessment of RNA half-lives.*

One million cells were plated in a single well of a 6-well plate and treated the next day with Flavopiridol (400 nM – S2679, Selleckchem) for the indicated times. All samples were harvested at the end of the assay for RNA extraction and RT-qPCR analysis.

*Oct4 knock-down.*

Oct4 knock-downs were performed with siRNAs (Dharmacon, ON-TARGETplus Mouse Pou5f1 siRNA – L-046256-00-0005) and compared to untergated control siRNAs (On Target plus control pool D-001810-10-05). Cells were nucleofected with 200 pmol of siRNA using a Mouse ES Cell Nucleofector kit (Lonza, VPH-1001, program A30) and cultured for 24h.

*Reverse transcription and real time polymerase chain reaction (RT-qPCR).*

Total RNAs were isolated with Trizol (Invitrogen, 15596026) according to the manufacture's protocol and absence of DNA contamination was ensured by additional DNase I digestion (Qiagen, 79254). Reverse transcription (RT) reactions were performed with random hexamers on 0,5 – 2 µg of total RNA according to manufacturer's instructions (First Strand cDNA Kit, Roche, 04379012001). Real-time PCR reactions were performed in duplicate in 384-well plates with a 480 Light Cycler (Roche) using Light Cycler 480 SYBR Green I Master Mix (Roche, 04707516001) and qPCR primers at 0.4µM final concentration. qPCR primers sequences are listed in Table S2. Standard and melting curves were generated to verify the amplification efficiencies (> 85%) and the production of single amplicons. Relative DNA amount was obtained from Cp (Crossing point) calculated from the second derivative of the DNA amplification signal over time. For RT-qPCR experiments, values for gene expression were normalized to the levels of housekeeping gene *Tbp* mRNA. Assays to measure RNA half-lives were normalized to the levels of *28s* mRNA.

*RNA-seq analysis and annotation of Suv39h1as.*

Poly-A selected RNAs were sequenced by Novogene Ltd (stranded, PE150) and reads aligned to the mm10 genome using STAR[44], quantified by RSEM[45] and counts transformed into Transcripts per million (TPM, Table S1). To annotate *Suv39h1as*, reads were mapped to the genome using Hisat2[46]. Bam files were used to build new transcript models with Stringtie[47] based on the Gencode (vM12) gtf (default parameters except -m 300). The resulting gtf files were merged (default parameters, except -m 300 -c 0.5 -F 0.5 -f 0.05), and all *Suv39h1as* isoforms annotated. Only those experimentally validated by Topo-cloning of poly-A selected cDNAs and RT-qPCR quantifications were retained. *Suv39h1as* isoforms 1 and 2, as reported in Fig.1, were validated by Topo-cloning and sequenced with primers Suv-as_ex1a-1-F/Suv-as_ex4-3-R and Suv-as_ex4s-1-R, respectively. Isoform 3 was validated with primers Suv-as_ex1b2-F/Suv-as_ex4-3-R. Splicing events were also validated by RT-qPCR: splicing from promoter 1a (Isoforms 1 and 2) and promoter 1b (Isoform 3) with exon 2, with primers Suv-as_ex1b2-F/Suv-as_ex1b2-R and Suv-as_ex1a2-F/Suv-as_ex1a2-R, respectively, and splicing from exon 2 and 3 with primers Suv-as_ex23-F/Suv-as_ex23-R. All primer sequences are available in Table S2.

*Chromatin immunoprecipitation (ChIP).*

Ten million cells were crosslinked either for 45 min with DSG 1X plus 10 min in Formaldehyde (FA) 1% in PBS 1X for TF binding analysis or only for 10 min in FA 1% for histone modification analysis. Formaldehyde was quenched with 125 mM glycine for 5 min at room temperature. Nuclei were prepared in 1ml of ice-cold swelling buffer (25 mM Hepes pH7.95, 10 mM KCl, 10 mM EDTA) freshly supplemented with 1× protease inhibitor cocktail (PIC-Roche, Cat# 04 693 116 001) and 0.5% IGEPAL (Sigma, Cat#I8896) for 20 min on ice and 50 passes in a dounce homogenizer. Nuclei were then resuspended in ice-cold D3 buffer (0.1% SDS, 15 mM Tris pH 7.6, 1 mM EDTA), freshly supplemented with 1×PIC, and sonicated using a Covaris M220-Setpoint at 6°C and 10 (FA only) to 15 (DSG/FA) cycles with the following parameters for each cycle: 60 sec duration, peak power of 67W, duty factor of 15% and cycles/burst of 500. A delay of 45 sec is added at the end of each cycle. Result of the average power is 10W. After centrifugation (15 min, 14000 rpm, 4 °C), the supernatant was stored at −80 °C until use. 20 µl were used to quantify the chromatin concentration and check DNA size (typically 200-600 bp). Fifteen to 20 µg of chromatin were used for each ChIP after pre-clearing it for 1.5 hours rotating on-wheel at 4 °C in 1 ml of TSE150 (0.1% SDS, 1% Triton X-100, 2 mM EDTA, 20 mM Tris-HCl pH8, 150 mM NaCl) buffer containing 50 µl of protein G Sepharose beads (Active Motif, Cat#37499) 50% slurry, previously blocked with BSA (0.5 mg/ml; Roche, Cat# 10711454001) and yeast tRNA (1 µg/ml; Roche Cat# 10109495001). Immunoprecipitations were performed overnight rotating on-wheel at 4 °C in 500 µl of TSE150. 20 µl were set apart for input DNA extraction and precipitation. 50µl of blocked protein G beads 50% slurry was added for 2h rotating on-wheel at 4 °C. Beads were pelleted and washed for 5 min rotating on-wheel at room temperature with 1 ml of buffer in the following order: 2× TSE150, 1 × TSE500 (as TSE150 but 500 mM NaCl), 1× washing buffer (10 mM Tris-HCl pH8, 0.25M LiCl, 0.5% IGEPAL, 0.5% Na-deoxycholate, 1 mM EDTA), and 2 × TE (10 mM Tris-HCl pH8, 1 mM EDTA). Elution was performed in 100 µl of elution buffer (1% SDS, 10 mM EDTA, 50 mM Tris-HCl pH 8) for 15 min at 65 °C after vigorous vortexing. Eluates were collected after centrifugation and beads rinsed in 150 µl of TE-1%SDS. After centrifugation, the supernatant was pooled with the corresponding first eluate. For both immunoprecipitated and input chromatin, the crosslinking was reversed overnight at 65 °C, followed by proteinase K treatment, phenol/chloroform extraction and ethanol precipitation._Input and IP samples were analyzed by qPCR using primers in Table S2. The 2dCt method was used. All values were corrected to the input. The antibodies used and their working dilution are indicated in Table S2.

*RNA/DNA Fluorescent In Situ Hybridization (FISH).*

Using the online tool “Stellaris RNA FISH probes designer” from LGB biosearch laboratories, single-strand probes for *Suv39h1* (47 oligos, 30 in exons, see sequences Table S2) and *Suv39h1as* (35 exonic oligos, see sequences Table S2) were designed for single-molecule FISH (smFISH). DNA probes for DNA-FISH were generated by nick translation (Vysis Nick Translation Kit; Abbott, cat. 32-801300) using a fosmid clone (WIBR1-2188H11 – from Children’s Hospital Oakland Research Institute, bacpac.chori.org) covering the entire locus. Cells were fixed in Formaldehyde 4%, quenched with Glycine (1M) and cytospun (Cytospin3, Shandon, at 400 rpm for 5 min with a low acceleration) onto slides that were kept in Ethanol 70% at 4°C until use. Slides were washed in 100% ethanol for 2 min and air dried. For each spot, 10 µL of hybridization cocktail (SSC2X – S6639-1L, Sigma; Dextran – Life technologies, 5%; Formamide

10% – Sigma F9037; 2 µg/µL E. Coli tRNAs – Sigma 10109541001; 5 mmol/L Ribonucleoside Vanadyl Complex – NEB S1402S; 0,5 µg/µL BSA – NEB B9001S) and smFISH probes for *Suv39h1* and *Suv39h1as* (each at 0,6 µmol/L) were used for overnight incubation into a humid chamber at 37°C. Slides were washed in fresh SSC 2X/Formamide 10% for 30 min at 37°C, mounted and counterstained with Vectashield Antifade Mounting medium with DAPI (Vector Laboratories, H-1200-10). Sm-FISH images were acquired with an inverted Nikon Eclipse X microscope equipped with: X63 oil immersion objective (N.A1.4); LUMENCOR excitation diodes; Hamamatsu ORCA-Flash 4.0LT camera; NIS Elements 4.3 software. The position of each image was recorded on the microscope. Subsequently, the coverslips were removed and the slides washed 3 times in washing medium (4X SSC, 0,2% Tween-20) at 37°C, and treated with RNaseA 10U/ml (Invitrogen, cat. EN0531) in 2XSSC at 37°C for 1h. DNA denaturation was performed in 50% formamide/2XSSC at 80°C for 30min. Slides were dehydrated in cold ethanol and hybridized overnight in a 50% Formamide/ 2X Hybridization cocktail (4X SSC – Sigma, S6639-1L; 20% Dextran sulfate – Life technologies; 2 mg/mL Bovine Serum Albumine –NEB B9001S; and 40 mM Ribonucleoside Vanadyl Complex– NEB S1402S) at 37°C with 0.3 ng of DNA-Fish probe, 3ul of mouse Cot1 DNA (Invitrogen, cat 18440016) and salmon sperm DNA (Invitrogen, cat.15632011), previously denatured in Formamide (7' at 75°C). After overnight hybridization of the probes, the slides were washed 3 times in 50% Formamide/2X SSC buffer at 37 °C for 5 min and 3 times in 2XSSC buffer at 37 °C for 5 min, mounted with Vectashield containing DAPI and imaging on the previous recorded positions.

*Western-blot.*

Cell were lysed in in Laemmli buffer (1 000 000 cells per µL ; #1610747, BioRad) at 95°C for 5 min and samples run in a mini-PROTEAN® TGX Stain-Free Precast Gel (Bio-Rad, 456-8086) in 25 mM Tris, 0,21 M Glycine, 50% SDS at 120V with constant voltage and transferred onto nitrocellulose membranes (Life Science, 10600003) in a 25mM Tris, 0.21 M glycine, 20% ethanol solution. The membrane was blocked in phosphate-buffered saline (PBS; 0.8 g/L NaCl, 0,02 g/L KCl, 0.144 g/L $Na_2HPO_4$, 0.024 g/L $KH_2PO_4$, pH = 7.2) with 0.1% Tween (PBST), 5% Bovin Serum Albumin (BSA) for 1 h and incubated in 3 ml of PBST 5% BSA with different antibodies listed Table S2, overnight at 4°C. Membranes were washed 3 times for 5 min in PBST and incubated with Pierce® goat anti-rabbit IgG-HRP conjugated secondary antibody (Thermo Scientific #314666, 0,1 µg/ml or 50 ng/ml). Membranes were washed in PBST and developed using a Pierce® ECL Western Blotting substrate kit (Thermo Scientific, #32109) or Pierce® ECL plus Western Blotting substrate kit (Thermo Scientific, #32134) for 5 min or 1 min at RT and luminescence detected using a ChemiDoc MP Imaging Systems with Image LabTMTouch Software Version 2.2.0.08.

*Immunostainings.*

Cells were trypsinized, counted and resuspended at 1 million/ml in FCS free medium (DMEM-Glutamax/100 mM 2-mercaptoethanol/NEAA 1X) into sterile 1.5 ml Eppendorf tubes. To ensure direct comparison of wild-type and mutant cells, they were then individually incubated either with 10 µM Rhodamine Red dye (InVitrogen, Cat#CMTPX C34552) or 1µM Deep Red dye (InVitrogen , Cat#C34565) for 20 to 40 min at 37°C. The labeled cells were then collected by centrifugation, washed with PBS1X, resuspended in DMEM/10%FCS+LIF medium and mixed at a 1:1 ratio for Rhodamine and Deep Red labelled cells (usually ~0.4M each). 800 000 mixed cells were seeded onto Poly-L-Ornithine/Laminin coated single wells of a µ-slide 4 well Ph+ibiTreat (Ibidi GmbH Ref#80446) and incubated for 6H at 37°C and 7% CO2. Cells were then fixed directly into the well with freshly prepared PFA 4% (Fisher Scientific, Cat#16431755) for 10 min at room temperature in the dark and washed twice in PBS1X for 10 min. Cells were permeabilized with PBS1X/0.1% Triton X-100 (Sigma, Cat#T8787) for 10 min at room temperature. After three washes with PBS 1X, cells were blocked with PBS 1X/3% Donkey Serum (Sigma, Cat#D9663) for 30 min in the dark and incubated overnight with primary antibodies (diluted in PBS 1X/10% DS). Following three washes with PBS, 1h incubation with secondary antibodies at room temperature in the dark and 3 washes with PBS 1X, nuclei were counterstained with DAPI (Sigma, Cat#D9542), washed in PBS1X and imaged with an inverted Nikon Eclipse X microscope equipped with: X20/0.45 (WD 8.2-6.9) objective; LUMENCOR excitation diodes; Hamamatsu ORCA-Flash 4.0LT camera; NIS Elements 4.3 software. Quantifications were performed using Cell Profiler[48]. For each experiment, Rhodamine/Deep Red labelled cells were attributed using the FlowJo software. For each experiment, the fluorescence intensity of mutant cells was normalised to the median of the corresponding wild-type cells intensities imaged on the same spot. The data was plotted using the ggplot2 package[49] in R.

*Statistical analyses*

Statistical significance of gene expression and chromatin immunoprecipitation differences measured by qPCR were assessed with Student's t-test. For the analysis of the locus-wide effects of the loss of *Suv39h1as* on H3K4me1 and me2, all values obtained with primer pairs located between coordinates +6 and +15 were considered; for the effects measured at the promoter region, the position showing the highest difference for each histone mark was used. Single molecule FISH data on RNA counts per cell was evaluated with Mann-Whitney-Wilcoxon tests and differences in transcription frequency with a Chi-squared test. Population differences observed with immunofluorescence were assessed with Kolmogorov-Smirnov tests. Differences in clonal assays were evaluated with Mann-Whitney-Wilcoxon tests using raw colony counts.
